# Quantifying epigenetic modulation of nucleosome breathing by high-throughput AFM imaging

**DOI:** 10.1101/2021.07.29.454136

**Authors:** Sebastian F. Konrad, Willem Vanderlinden, Jan Lipfert

## Abstract

Nucleosomes are the basic units of chromatin and critical to the storage and expression of eukaryotic genomes. Chromatin accessibility and gene readout are heavily regulated by epigenetic marks of which post-translational modifications of histones play a key role. However, the mode of action and the structural implications on the single-molecule level of nucleosomes is often still poorly understood. Here, we apply a high-throughput AFM imaging and analysis pipeline to investigate the conformational landscape of the nucleosome variants H3K36me3, H3S10phos and H4K5/8/12/16ac. Our data set of >25,000 nucleosomes reveals nucleosomal unwrapping steps corresponding to 5 bp DNA. We find that H3K36me3 nucleosomes unwrap significantly more than wild type nucleosomes and additionally unwrap stochastically from both sides similar to CENP-A nucleosomes and in contrast to the highly anti-cooperative unwrapping of wild type nucleosomes. Nucleosomes with H3S10phos or H4K5/8/12/16ac modifications show unwrapping populations similar to wild type nucleosomes and also retain the same level of anti-cooperativity. Our findings help putting the mode of action of these modifications into context: While H3K36me3 likely partially acts by directly affecting nucleosome structure on the single-molecule level, H3S10phos and H4K5/8/12/16ac must predominantly act through higher-order processes. Our analysis pipeline is readily applicable to other nucleosome variants and will facilitate future high-resolution studies of the conformational landscape of nucleoprotein complexes.

**Statement of Significance:** The packing and readout of our genome is tightly regulated by post-translational histone modifications (PTMs). While a vast range of PTMs has been studied with respect to their implications for gene activity and replication, a detailed view of the direct effect of PTMs on conformational changes of nucleosomes is still lacking. Here we investigate the structural implications of several key modifications (H3K36me3, H3S10phos and H4K5/8/12/16ac) by high-throughput AFM imaging. Our findings enable a better understanding of the mode of action of these specific modifications and provide an analysis pipeline for the investigation of other epigenetic modifications.

---

Nucleosomes are the fundamental units of compaction of eukaryotic DNA into chromatin and function as regulators of gene readout and activity^1–3^. Canonical nucleosome core particles consist of two copies each of the four histones H2A, H2B, H3 and H4 assembled into a histone octamer that is wrapped by ~147 bp of DNA^4,5^. Electrostatic interactions and specific molecular contacts stably pack the DNA onto the histone octamer, yet DNA breathing, sliding, gaping, and loosening allow for nucleosomal dynamics on millisecond to minute time scales^6–8^.

Post-translational modifications (PTMs) of histones play a key role in the formation of higher order chromatin structure^3,9–13^, the recruitment of proteins and complexes with specific enzymatic activities^14^, and the maintenance of DNA repair^15^ and replication^16^. Numerous histone variants and PTMs alter histone-histone and histone-DNA interactions^17–19^ to yield nucleosomal structures with varying degrees of stability and DNA wrapping. Specifically, PTMs at the N-terminal tails of histones H3 and H4 located next to the DNA entry-exit sites can affect DNA opening dynamics by introducing additional charge, by neutralizing existing charge, or by adding steric constraints^2,20^. Among the astonishing number of PTMs^21,22^, the most frequent PTMs at the histone-DNA interface are methylations, acetylations and phosphorylations^2,14^. Acetylation neutralizes the positive charge of lysine and phosphorylation introduces negative charge. Methylation does not alter the charge of the histone protein but, similar to acetylation and phosphorylation, adds steric bulk to the system.

While many studies have investigated post-translational modifications (PTMs) with respect to their effects on nucleosomal structural dynamics^23,24^ and on the interaction with nucleosome- or DNA-binding proteins^25–27^, a detailed investigation of the effect of distinct PTMs on nucleosome wrapping is currently lacking. It is critical to understand the direct effects of PTMs on nucleosome conformations, as they can influence the accessibility of nucleosomal DNA for readout and processing and can modulate the conformational landscape that underlies interactions with additional binding partners.

Here, we use a high-throughput pipeline based on atomic force microscopy (AFM) imaging to investigate the conformational landscape of nucleosome variants with several key post-translational N-terminal tail modifications on histones H3 and H4: H3K36me3, H3S10phos and H4K5/8/12/16ac (Fig. 1a). These specific modifications are selected for several reasons: First, our goal is to investigate a range of different nucleosome modifications and, therefore, cover trimethylation, acetylation and phosphorylation. Second, we aim for modifications at different positions in the histones. While H3K36me3 and H4K5/8/12/16ac lie close to the DNA entry/exit region of histones H3 and H4 respectively, H3S10phos is located more distal towards the start of the N-terminal tail of histone H3. Third, for H3K36me3^28,29^ and H4K5/8/12/16ac^30^, previous measurements of the nucleosome structure found no direct effect of the modifications. Yet, due to the close proximity of both modifications to the DNA entry/exit site, we hypothesized that these PTMs could have an effect on the nucleosome wrapping landscape and aimed to detect it with our sensitive assay. Likewise, H3S10phos is an interesting modification as it is involved in both transcriptional activation and chromatin compaction^31^, two structurally opposed processes, therefore raising the question whether H3S10phos has structural implications on the nucleosome itself or merely acts as a protein binding platform.

**Figure 1.**
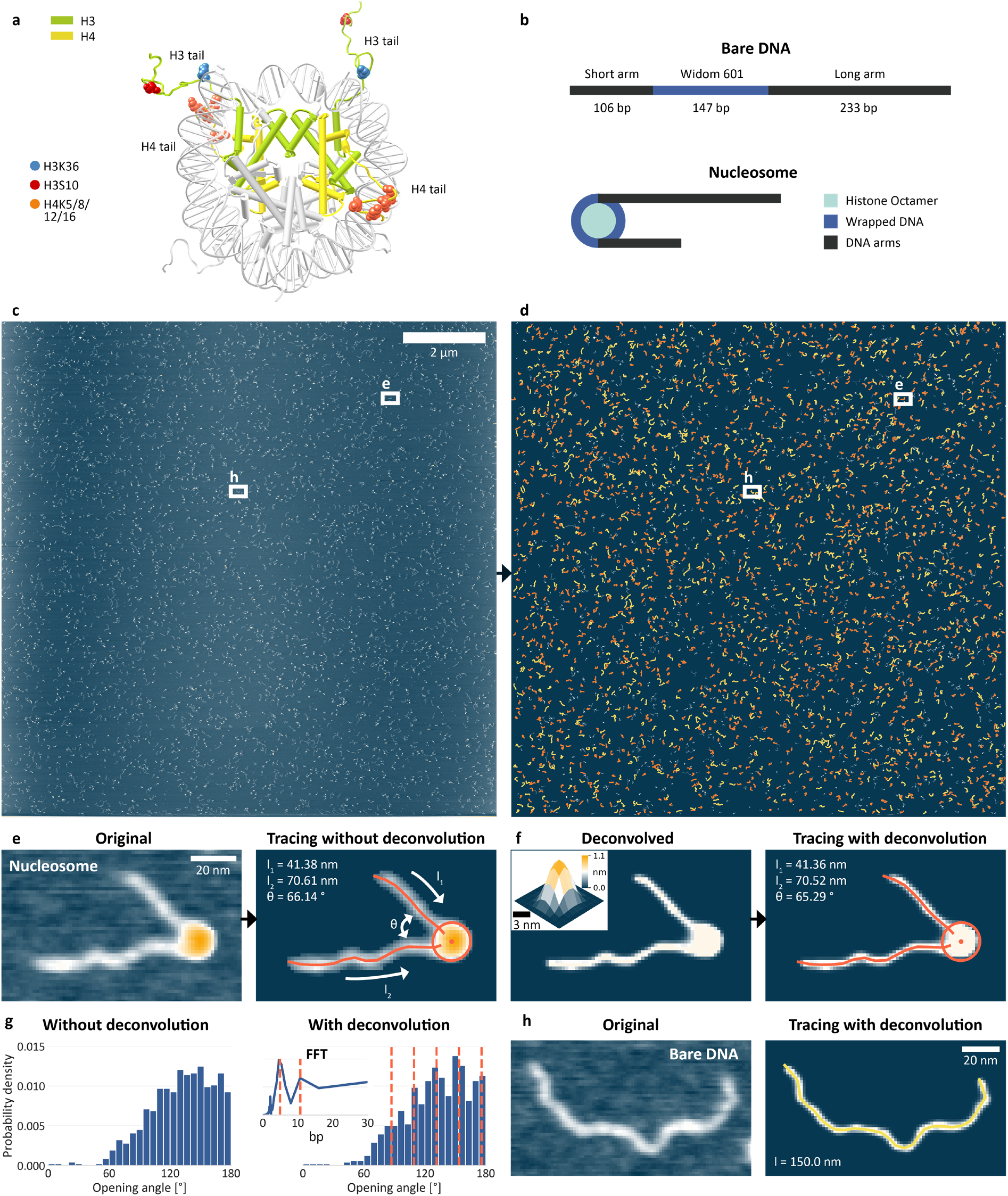
DNA and nucleosome structure parameters from automated AFM image analysis. **a,** Crystal structure of a canonical nucleosome (PDB 1KX5). Colored spheres represent the positions of the modified amino acids in the histone tail considered in this work. Among the three histone tail modifications investigated are H3K36me3 (three additional methyl groups at lysine 36 of histone H3 – blue spheres), H3S10phos (phosphorylation of H3 histones at serine 10 – red spheres) and H4K5/8/12/16ac (acetylation of H4 histones at lysines 5, 8, 12 and 16 – orange spheres) **b,** Schematic of the construct used throughout this work. The 486 bp DNA consists of a 147 bp W601 nucleosome positioning sequence that is flanked by a short and a long arm of 106 bp and 233 bp, respectively. Histone octamers contain two copies each of H2A, H2B, H3 and H4. **c,** AFM image of bare DNA and nucleosomes with a field of view of 12 μm x 12 μm at a resolution of 1.46 nm/pixel (8192^2^ pixels). **d,** Traces of 901 bare DNA strands (orange) and 1624 nucleosomes (yellow) obtained by the automated image analysis pipeline from the image shown in c. **e,** Zoom of a nucleosome image before and after tracing. The zoom area is indicated in panels c and d. **f,** Same nucleosome image as panel e after Richardson-Lucy deconvolution. The inset displays the shape of the AFM tip deduced from the bare DNA molecules in the same AFM image and used for deconvolution. **g,** Opening angle distribution for the same data set analyzed without and with deconvolution. The deconvolved data shows the 20° (5 bp) unwrapping periodicity of nucleosomes (N=716, only partially unwrapped nucleosomes shown). **h,** Bare DNA before and after tracing.

AFM imaging is a powerful tool to probe DNA and nucleosome structure^32–38^ and we have recently developed a multi-parameter image analysis pipeline to quantify the wrapping of nucleosomes with nanometer resolution, label-free, and at the single-molecule level^38^. Here, we have improved the resolution of our assay by adding a deconvolution step to allow for more accurate parameter tracing, enabling the direct observation of the nucleosomal unwrapping periodicity of 5bp from nucleosomal opening angles. We find nucleosomes with the H3K36me3 modification to occur significantly less likely in the fully wrapped state compared to canonical nucleosomes and to exhibit stochastic instead of anti-cooperative unwrapping. In contrast, H4K5/8/12/16ac and H3S10phos do not show significant changes in both unwrapping and anti-cooperativity compared to canonical nucleosomes. We discuss these results in the context of biological function and epigenetic regulation of genome organization.

## Results

### Quantifying nucleosome conformations *via* automated AFM image analysis with deconvolution

We assembled nucleosomes by salt gradient dialysis under sub-stoichiometric conditions, such that the final sample contains bare DNA and predominantly mono-nucleosomes. We use a 486 bp DNA construct that features a W601 nucleosome positioning sequence^39^ (147 bp) flanked by a short DNA arm (106 bp) and a long arm (233 bp) (Fig. 1b and Methods). We deposited samples from aqueous buffer on poly-L-lysine coated mica prior to rinsing and drying of the sample. High-resolution images of the deposited nucleosome samples were obtained by amplitude modulation AFM in air (Fig. 1c).

To quantify nucleosome conformations from the AFM images, we build on our previously published AFM image analysis pipeline to trace bare DNA and nucleosomes in the AFM images by multi-parameter analysis^38^ and extend it by adding an additional deconvolution step that allows for more accurate tracing. The tracing consists of two steps: First, bare DNA and nucleosomes are detected and classified by subtracting the background and consecutively utilizing the topology of the one pixel wide backbone – the skeleton – of the molecules (Supplementary Fig. 1). Second, structural parameters of the classified molecules are extracted by automatically tracing the molecules with the custom analysis software. The extracted parameters comprise contour length and bend angles for bare DNA and core particle height and volume, arm lengths, and opening angle for nucleosomes (Fig. 1e-h). The vectors connecting the ends of the arm entry/exit region of the core particle and the center of the core particle define the nucleosome opening angle. In particular, the combined information of free DNA contour length and opening angle enables to identify the unwrapping state of each nucleosome, i.e. to classify how the DNA wraps around the histone core.

To further increase the accuracy of our assay compared to previous applications of AFM imaging to nucleosome conformations, we implemented an image deconvolution step. In general, the dimensions of molecules are overestimated in AFM imaging due to the finite size of the AFM tip^38,40^. In particular, we find that tip convolution obscures the exact entry/exit position of DNA in the nucleosome images. We estimate the shape of the AFM cantilever tip (Fig. 1f inset, Supplementary Fig. 2 and Methods) from the bare DNA in our images and typically find tip shapes with an end radius of 5-6 nm (Fig. 1f inset) in line with the size as specified by the manufacturer (see Methods). We use this tip shape estimate for subsequent image deconvolution based on the Richardson-Lucy algorithm^41,42^ (see Methods). Applying the tip deconvolution leads to sharper images, in particular evident from the DNA paths (Fig. 1f, h).

Comparing the opening angles measured with and without image deconvolution, demonstrates the considerable impact of this approach (Fig. 1g). While the angle distribution of nucleosomes traced without deconvolution gives a broad and relatively featureless distribution of opening angles, the distribution of opening angles traced after applying the deconvolution clearly indicates a periodicity in the opening angle distribution of ~20°, i.e. 5 bp of unwrapping (Fig. 1g, inset – a fully wrapped nucleosome wraps 147 bp in ~1.7 turns). This 5 bp unwrapping periodicity ultimately stems from the periodicity of the DNA helix and is in line with results from single-molecule DNA force spectroscopy experiments^43,44^ and with cryo-EM observations of nucleosome wrapping states^45^.

### Quantifying nucleosome wrapping populations by multi-parameter analysis

To quantify the length of DNA wrapped in the nucleosomes from AFM data, we evaluated the average contour length of the bare DNA molecules in each image (Supplementary Fig. 3) and similarly measured the nucleosome arm lengths. By subtracting the combined arm lengths of individual nucleosomes from the mean contour length of bare DNA molecules, we obtain the wrapped length, i.e. the length of DNA confined in the nucleosome core particle. Simultaneously, we obtain the opening angle between the DNA segments entering the nucleosome particle for each nucleosome. The 2D distribution of nucleosome opening angles and nucleosome wrapping provides a quantitative view of the nucleosome wrapping landscape^38^ (Fig. 2). For a representative data set of canonical nucleosomes, the 2D kernel density distribution reveals two major populations (Fig. 2a): One that features wrapped lengths >150 bp and opening angles <100° and one that features wrapped lengths <150 bp and opening angles >70°. We have previously identified^38^ the population of nucleosomes with wrapped lengths <150 bp as partially unwrapped. This population features a negative correlation between opening angle and wrapped length, since the opening angle increases by further unwrapping of the DNA arms. Similarly, we have previously assigned the remaining population with wrapping of >150 bp of DNA to fully wrapped nucleosomes. Previous simulations of nucleosomes in AFM imaging^38^ rationalize why the apparent wrapped lengths for fully wrapped nucleosomes exceed the 147 bp expected from the crystal structure: the DNA arms that leave the nucleosome entry/exit site overlap close to the nucleosome core particle. In the images, the crossing DNA strands lead to an underestimation of the length of the DNA arms, resulting in longer apparent wrapped lengths for fully wrapped nucleosomes. Utilizing the local minimum seen in the principal component analysis (PCA) of nucleosome core volumes and opening angles (Fig. 2a, inset), we separated fully wrapped and partially unwrapped nucleosomes (white and black dots respectively) and find that in this particular data set of unmodified nucleosomes, 31% of the nucleosomes are fully wrapped and 69% of the nucleosomes are partially unwrapped.

**Figure 2.**
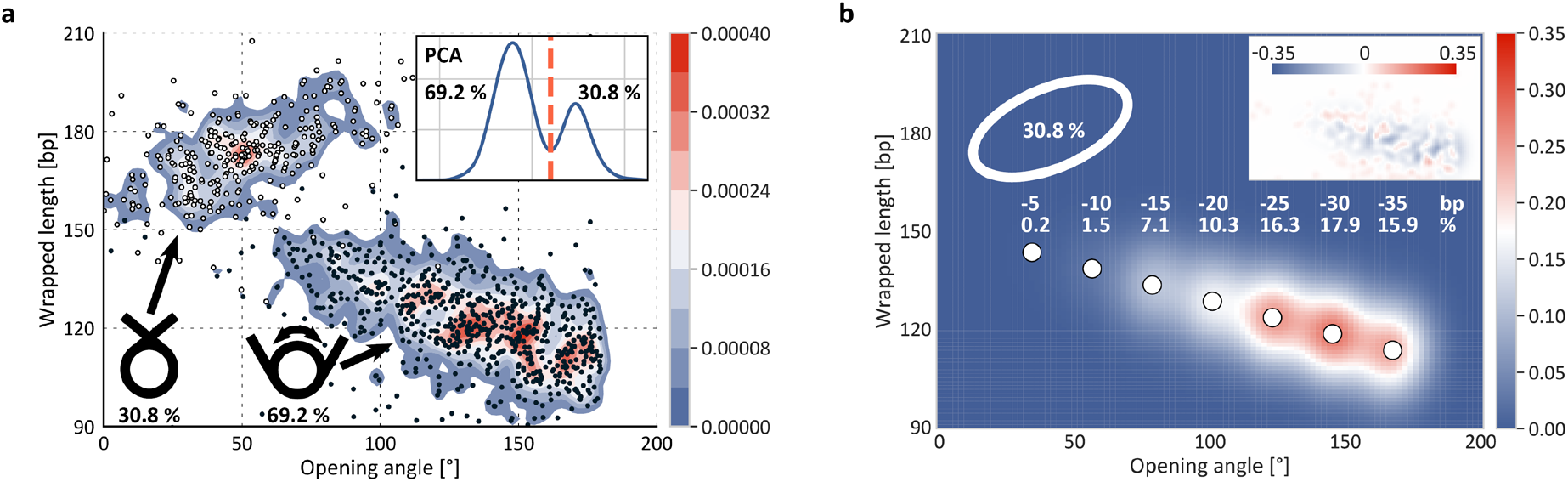
Estimating nucleosome wrapping populations. **a,** Wrapped length versus opening angle distribution for canonical nucleosomes. White and black circles represent individual nucleosomes (N = 1035), the colored contours are the 2D kernel density estimate. The inset shows a principal component analysis (PCA) of wrapped length and volume that is used to separate the two nucleosome populations (fully vs. partially wrapped). **b,** 2D Gaussians fit to the density distribution of the partially unwrapped nucleosomes. The Gaussian amplitudes represent the populations of the 5 bp unwrapping substates; the inset shows the residuals of the fit.

To quantitatively investigate nucleosome unwrapping, we fitted the distribution of partially unwrapped nucleosomes with seven 2D Gaussians–one Gaussian per 5 bp unwrapping step up to an unwrapping of 35 bp–located at fixed distances and corresponding to the 5 bp unwrapping periodicity (Fig. 2b). The amplitudes of the Gaussians represent the occupancies of the individual states of unwrapping and show that for unmodified nucleosomes most of the partially unwrapped nucleosomes unwrap 25 to 35 bp of DNA. To quantify how reproducibly our analysis pipeline can determine the wrapping populations, we performed independent repeat measurements–all from independent nucleosome reconstitutions and two protein batches– and applied the same analysis pipeline to the separate data sets to obtain mean wrapping distributions and errors. Each repeat comprises >1000 individual nucleosome molecules. Our method is highly reproducible and yields very precise estimates of the individual wrapping populations: the average absolute SEM for the populations is ~1 %. Therefore, our analysis pipeline provides a highly accurate and quantitative assay to investigate the effect of epigenetic modifications on nucleosome structure.

### Post-translational modifications alter wrapping of H3K36me3 nucleosomes

To study how post-translational histone tail modifications affect nucleosome wrapping on the single-molecule level, we applied our analysis pipeline to two nucleosome constructs that have PTMs close to the DNA entry/exit region, H3K36me3 and H4K5/8/12/16ac, and one nucleosome construct that has a PTM further outside of the nucleosome core particle at the end of the histone H3 tails: H3S10phos (Fig. 3). As for the unmodified nucleosomes, we performed 4-5 independent repeat measurements from independent nucleosome reconstitutions and two protein batches all measured on different days for each nucleosome variant.

**Figure 3.**
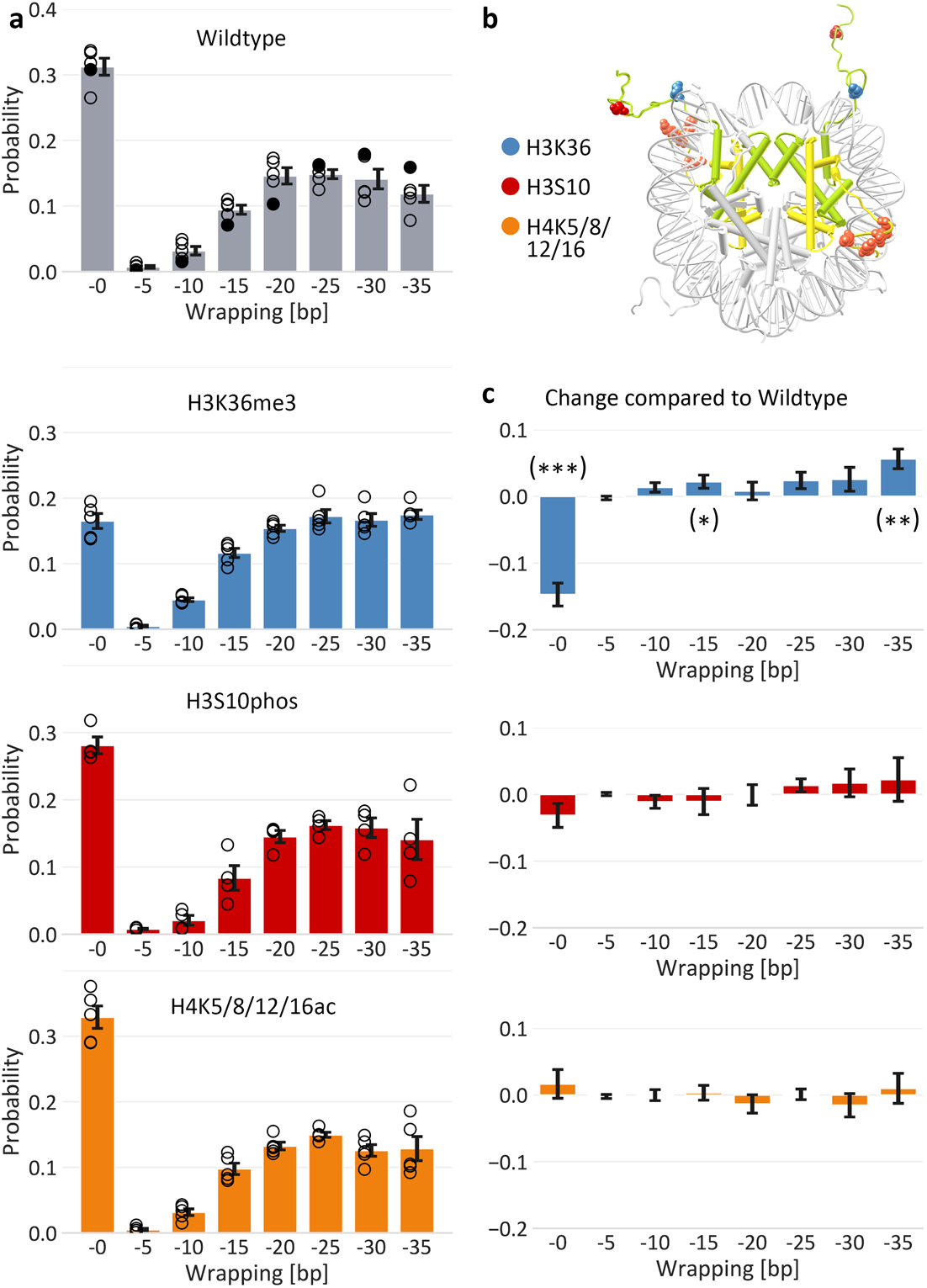
DNA wrapping populations of post-translationally modified nucleosomes. **a,** Populations of DNA wrapping conformations for unmodified nucleosomes and the three modified nucleosome constructs containing either three methylations at histone H3 lysine 36, a phosphorylation at histone H3 serine 10 or acetylations at histone H4 lysines 5, 8, 12 and 16. The populations were determined from high-throughput analysis of AFM images as shown in Fig. 2; the filled black circles in panel a are from the data set in Fig. 2. For each histone variant four to five independent measurement repeats were obtained; circles indicate the populations of the individual data sets; bars and error bars are the mean and SEM from the independent repeats. **b,** Crystal structure of the canonical nucleosome (PDB 1KX5). Colored spheres represent the positions of the modified histone tail amino acids. **c,** Differences between the wrapping populations of the modified nucleosomes and the unmodified nucleosomes. Significant differences–as determined from two-sample t-tests–are indicated by stars.

For H3K36me3 nucleosomes, *i.e*. nucleosomes with three methyl moieties on the epsilon amino group of lysine residue 36 of the histone H3 tails, we find that only a small fraction populates the fully wrapped state (16.5 % ± 1.1 %; mean + SEM from five biological repeats compared to 31.2 % ± 1.3 % for canonical nucleosomes, Fig. 3a) and the vast majority of nucleosomes populates states of partial unwrapping (83.5 % ± 1.1 %). H3K36me3 nucleosomes are almost two fold less likely to occupy the fully wrapped state compared to canonical nucleosomes and are significantly more likely (*p* = 0.005 from a two-sample t-test) to populate states of higher unwrapping at −35bp (Fig. 3c). Thus, trimethylation at H3K36 alters nucleosome structure towards increased unwrapping. Previous FRET studies did not find a measurable difference in nucleosome unwrapping between H3K36me3 and canonical nucleosomes^28,29^. However, these studies rely on the binding of the repressor protein LexA to the partially unwrapped nucleosomes between base pairs 8 and 27 and thus require a change in the states of unwrapping >25 bp to detect differences between H3K36me3 and canonical nucleosomes. Our data show that only part of the additionally unwrapped H3K36me3 nucleosomes populate these states of higher unwrapping and thus the overall increased unwrapping might not be detected by the FRET/LexA methodology at the same level of detail as in our assay.

For H3S10phos nucleosomes, *i.e*. nucleosomes with phosphorylated histone tails at serine residue 10, we find no significant differences in partial unwrapping compared to canonical nucleosomes (two-sample t-test *p* = 0.13). 28.1 % ± 1.3 % of the phosphorylated nucleosomes are fully wrapped and 71.9 % ± 1.3 % are partially unwrapped. Phosphorylation introduces negative charge to the serine and thus affects the electrostatic potential of the N-terminal histone tail. However, the modified serine lies on the outer end of the histone tail (Fig. 3b) and therefore H3S10phos appears to have only a small effect on the intrinsic nucleosome structure, in line with a previous study that suggested that H3S10phos does not merely act by creating an open chromatin configuration in which DNA is more accessible to the transcriptional machinery^46,47^.

For H4K5/8/12/16ac nucleosomes, *i.e*. nucleosomes with acetylated histone H4 tails at lysine residues 5, 8, 12 and 16, we find 32.9 % ± 1.7 % of the nucleosomes to occupy the fully wrapped state. 67.1 % ± 1.7 % occupy states of partial unwrapping with most of them unwrapping 20 to 35 bp of DNA (Fig. 3a), corresponding to no significant differences in wrapping between the tetraacetylated and canonical nucleosomes (*p* = 0.46; Fig. 3c). Histone tail acetylations neutralize the positive charge of the modified lysines and thus reduce electrostatic interactions between the histone tails and the negatively charged DNA. Our observation for H4K5/8/12/16ac nucleosomes is in agreement with a recent single-molecule study that found no increased unwrapping for nucleosomes that contained 12 – so three-fold more than in our construct – H4 tail lysine acetylation mimics^48^. Similarly, a FRET study found no effect of H4 acetylations on DNA entry/exit site geometry at ionic conditions same as ours^49^. We speculate that hydrogen bonding and hydrophobic forces outweigh electrostatic interactions in the binding between histone H4 tail lysines 5, 8, 12 and 16 and DNA as proposed for H4K16 in a previous simulation study on the effect of H4K16ac on the histone-DNA binding affinity^50^.

### Post-translational modifications can affect nucleosome unwrapping pathways

Previous studies based on single-molecule manipulation, FRET^51^, and cryo-EM^45^ revealed that unwrapping at one exit site stabilizes binding at the second exit site, leading the anti-cooperative unwrapping of DNA from nucleosomes. We have recently shown that by analyzing the distribution of short arm lengths *vs*. opening angles, our high-throughput AFM image analysis approach is sensitive enough to detect this anti-cooperative unwrapping of canonical nucleosomes^38^. In our data, the anti-cooperative opening of nucleosomes becomes apparent for the nucleosomes at opening angles >80°, *i.e*. in the regime of partially unwrapped nucleosomes (Fig. 4a). The distribution of partially unwrapped nucleosomes splits into two branches reflecting the anti-cooperative nature of the unwrapping process^38^.

**Figure 4.**
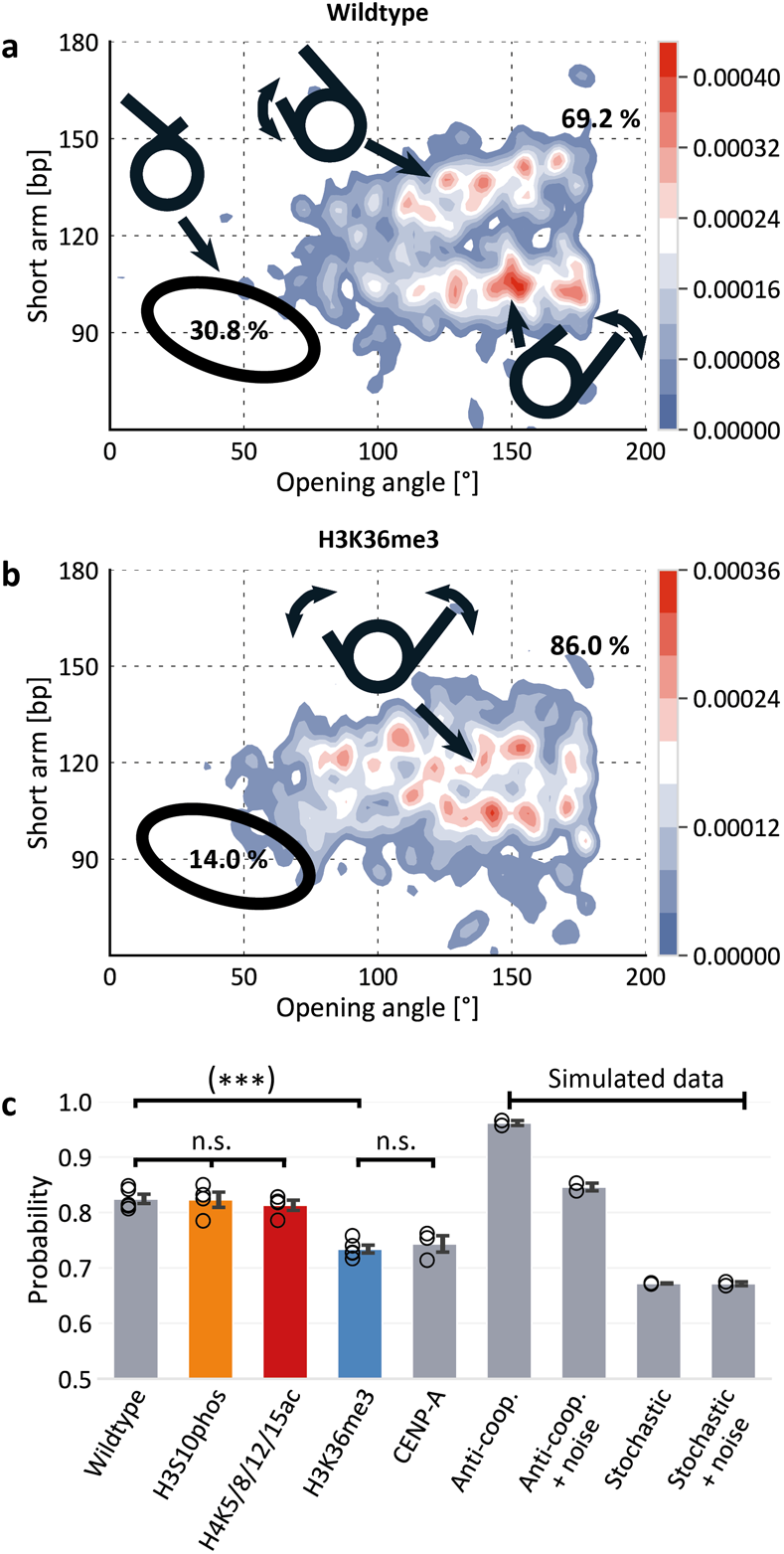
Unwrapping pathways of post-translationally modified nucleosomes. **a,** 2D kernel density profile (bandwidth = 2.5°, 2.5 bp) of short arm length and opening angle for H3 nucleosomes. A bimodal distribution for opening angles >80° is apparent, consistent with anti-cooperative unwrapping of the nucleosome core particle (*N* = 1035). The distribution of fully wrapped nucleosomes (30.8 % of all nucleosomes, indicated by the black ellipse) was omitted from the plot for clarity. **b**, 2D kernel density profile (bandwidth = 2.5°, 2.5 bp) of short arm length and opening angle for H3K36me3 nucleosomes (*N* = 1155). **c,** Quantification of the tendency of the different epigenetically modified nucleosomes to unwrap anti-cooperatively or not (Supplementary Fig. 4). Unmodified, H3S10phos, and H4K5/8/12/15ac nucleosomes show similar high levels of anti-cooperative unwrapping; in contrast, H3K36me3 and CENP-A nucleosomes unwrap less anti-cooperatively.

To investigate the effect of epigenetic modifications on the cooperativity of nucleosome unwrapping, we calculate the probability of a certain nucleosome type to unwrap anti-cooperatively (Supplementary Fig. 4). For this purpose, we define an area in the 2D opening angle *vs*. short arm length density distribution (Fig. 4a and Supplementary Fig. 4) in which the nucleosomes are expected to lie in the case of anti-cooperative unwrapping and compare the population size to the fraction of nucleosomes outside that area. For the canonical nucleosomes, 82.5 % ± 0.8 % (mean + SEM from five biological repeats) are in the anti-cooperative unwrapping regime. Similarly, the H3S10phos and the H4K5/8/12/16ac nucleosomes occupy the anti-cooperative unwrapping regime at 82.3 % ± 1.4 % and at 81.3 % ± 0.9 % respectively (Fig. 4c), indicating that these modifications do not affect the anti-cooperativity in nucleosome unwrapping.

In contrast, we find a significant reduction in anti-cooperativity for the unwrapping of H3K36me3 (73.4 % ± 0.7 %) compared to the canonical nucleosomes (two-sample t-test *p* = 3.5o10^-5^), implying that a substantial part of H3K36me3 unwraps stochastically from both sides. Previously, we have observed a similar effect^38^ for nucleosomes that contained the histone H3 variant CENP-A (Fig. 4c). We have speculated that the shortened N-terminal alpha helix (αN) of CENP-A nucleosomes –compared to the larger H3 αN of canonical nucleosomes-might cause the loss of anti-cooperativity, in line with a previous cryo-EM study that has suggested that allosteric changes involving H3 αN might invoke anti-cooperative unwrapping in canonical nucleosomes^45^. Comparing our findings for H3K36me3 nucleosomes to the CENP-A data from the previous study shows similar reduction of anticooperativity (Fig. 4b) for both nucleosome types. Both exchange of H3 with CENP-A and trimethylation at H3K36 introduce changes to the histone octamer at the entry/exit region of the DNA and a reduced fraction of fully wrapped nucleosomes. Our finding of reduced anti-cooperativity in the unwrapping of H3K36me3 nucleosomes indicates that already subtle changes at the DNA entry/exit site of nucleosomes can strongly affect nucleosomal dynamics and opening pathways. To check whether differences in nucleosome positioning along the W601 sequence between unmodified and H3K36me3 nucleosomes plays a role in the different density distribution of H3K36me3 nucleosomes, we compared the positioning of both unmodified and trimethylated nucleosomes but found no differences (Supplementary Fig. 5).

To further understand the nature and extent of anti-cooperative unwrapping, we simulated synthetic AFM images of nucleosomes that explored two extreme scenarios: either exhibiting only anti-cooperative unwrapping or completely stochastic unwrapping (Supplementary Fig. 6). In short, we placed a disk, representing the nucleosome core particle, on a surface and simulated protruding DNA arms with different lengths at opening angles as deduced from the unwrapping state and the nucleosome crystal structure. The populations of unwrapping simulated are based on the experimentally determined unwrapping populations for unmodified nucleosomes (see Methods and Supplementary Fig. 6 for more detail).

Applying our analysis pipeline to simulated nucleosome images that exhibit completely anti-cooperative unwrapping and contain no added noise, we find very high scores for anti-cooperative unwrapping, >95%, as expected. This value is higher than what we observe for any of the experimental conditions. However, if we add Gaussian noise with a width of 5 bp, corresponding to approximately one pixel in our AFM images and representative of our imaging noise, to the short arm length, we find anti-cooperativity values of 84 % ± 0.7%, which are still slightly higher, but close to the experimentally observed values for canonical, H3S10phos and H4K5/8/12/16ac nucleosomes, suggesting that our data are consistent with these types of nucleosomes exhibiting almost perfectly anti-cooperative unwrapping. Conversely, if we simulate nucleosomes that unwrap randomly from either site, we find anticooperativity scores of 67 % ± 0.3 %, essentially independent of whether noise is added to the images or not due to the already stochastic nature of the distribution (Supplementary Fig. 6). The anti-cooperativity scores for the randomly unwrapping simulations are lower than any of the experimentally determined values, but relatively close to values determined for H3K36me3 and CENP-A nucleosome, suggesting that while H3K36me3 and CENP-A unwrap mostly random, they appear to retain some anti-cooperativity.

## Conclusions

Quantitative assessment of conformations of post-translationally modified nucleosomes is a key to understanding the mode of operation of the histone code. PTMs can have manifold effects on chromatin structure such as entry site unwrapping, nucleosome destabilization, chromatin fiber destabilization, and histone-histone destabilization^2,17,22^. In this work, we utilized a high-throughput image analysis pipeline to study the effect of the post-translational modifications H3S10phos, H3K36me3 and H4K5/8/12/16 on nucleosome structure and dynamics. From a multi-parameter analysis of >25,000 nucleosomes, we obtain a comprehensive and quantitative view of the molecular ensembles, which in turn allows us to extract detailed information about nucleosome wrapping with as little as 1 % uncertainty (SEM) for the populations of the individual unwrapping states.

The H3K36me3 modification exhibited the strongest effect on nucleosome wrapping probably due to its location at the DNA entry/exit site of the nucleosome. While we observe partial unwrapping of ~70% of the canonical nucleosomes, ~85% of H3K36me3 nucleosomes occupied states of partial unwrapping. Strikingly, in stark contrast to the anti-cooperative unwrapping of canonical nucleosomes where unwrapping from one side inhibits unwrapping from the other, H3K36me3 nucleosomes tend to unwrap stochastically from both sides. H3K36me3 acts via recruiting a number of histone PTM binding domains^27^ and is associated with DNA repair, alternative splicing, and transcription^2,52^. Work in drosophila suggests that H3K36me3 is enriched in gene bodies, in particular, in the region of transcribed genes distal to the transcription start site^53,54^. The increased proneness of H3K36me3 nucleosomes to partially unwrap suggests that H3K36me3 can directly affect higher order chromatin structure by increasing the heterogeneity of nucleosome-nucleosome contacts as well as the effective nucleosome valency^11^.

On the macromolecular level, histone tails play a key role in the formation of higher-order chromatin structures^3^. Acetylation of the histone tails inhibits the folding of the nucleosome array in vitro^9^ and elevated histone acetylations increase chromatin accessibility^55^ and reduce the clustering of nucleosomes^56^ in vivo. Additionally, H4 acetylation blocks the interaction between the H4 tail and the acidic patch of adjacent nucleosomes and thus decreases inter-nucleosomal interactions^57^. Yet, on the single-molecule level, we found no significant changes in nucleosome accessibility due to the H4K5/8/12/16ac modification.

Similarly, while we did not see significant changes in nucleosome wrapping for H3S10phos nucleosomes compared to canonical nucleosomes, it has previously been shown that, in principle, phosphorylation can have significant effects on nucleosome dynamics^58^. However, in these studies the phosphorylation occurred closer to the nucleosome dyad as compared to the phosphorylation investigated in our study that lies towards the end of the histone H3 tails. We speculate that H3S10phos predominantly acts by binding proteins such as certain members of the 14-3-3 family with H3S10phos specificity^59,60^ and also via cross-talk with other PTMs such as blocking of H3K9ac^61,62^ or promotion of H3K14ac^63^.

Our results highlight how different PTMs involved in transcriptionally active chromatin act through a range of mechanisms. We show that our high-throughput, high-resolution pipeline can reveal the effects of subtle chemical modifications on nucleosome conformations. More broadly, our approach is readily applicable to other nucleosome modifications and variants as well as their interactions with binding partners.

## Author contributions

All authors designed research. S.F.K. performed research, analyzed data, and wrote the manuscript with input from all authors.

## Acknowledgement

We thank Philipp Korber and Felix Müller-Planitz for help with initial nucleosome reconstitutions, Pauline Kolbeck, Tine Brouns, Wout Frederickx, Herlinde De Keersmaecker, Steven De Feyter, and Björn H. Menze for discussions and assistance with AFM imaging, and Thomas Nicolaus for help with sample preparation. This work was funded by the Deutsche Forschungsgemeinschaft (DFG, German Research Foundation) through SFB863 – Project ID 111166240 (A11).

## Methods

### DNA purification and nucleosome reconstitution

DNA was PCR amplified from a GeneArt High-Q String DNA fragment (Thermo Fisher Scientific, Waltham, Massachusetts) containing the Widom 601 positioning sequence. The DNA was purified using a QIAquick PCR purification kit (Qiagen, Hilden, Germany) and subsequently eluted to a volume of 30 μL in milliQ water. Unmodified and modified histone proteins were purchased from EpiCypher (Durham, North Carolina). Nucleosome reconstitutions were performed by salt gradient dialysis^64^. The dialysis chambers contained 0.65 μg of the histone octamers and 3 μg of the 486 bp DNA at 2 M NaCl and were placed in one liter of high-salt buffer (2 M NaCl, 10 mM Tris, 1 mM EDTA). Over the course of 15 hours, three liters of low-salt buffer (50 mM NaCl, 10 mM Tris, 1 mM EDTA) were transferred to the high-salt buffer at 4°C. Finally, the dialysis chambers were moved to one liter of low-salt buffer for three hours.

### AFM sample preparation and imaging

Poly-L-lysine coated mica was prepared by depositing 20 μL poly-L-lysine (0.01% w/v) on freshly cleaved muscovite mica for 30 seconds and subsequently rinsing the surface with 50 mL of milliQ water before drying with a gentle stream of filtered N_2_ gas. A sample mix containing bare DNA and reconstituted nucleosomes –usually 30% to 50% of the DNA strands do not bind to histones–was incubated at 200 mM NaCl and 10 mM Tris-HCl, pH 7.6, for all measurements for 1 min on ice. The sample mix is then deposited on the poly-L-lysine coated muscovite mica for 30 seconds and subsequently rinsed with 20 mL milliQ water before drying with a gentle stream of filtered N_2_ gas.

We used two different commercial AFM instruments for imaging. All AFM images were acquired in tapping mode at room temperature. One set of images was acquired on a Multimode VIII AFM (Bruker) using silicon tips (AC160TS, drive frequency of 300-350 kHz, tip radius 7 nm, Olympus, Tokyo, Japan). Images were scanned over a field of view of 3 μm x 3 μm at 2048 x 2048 pixels with a scanning speed of 1 Hz. Independent measurement repeats were performed on a Nanowizard Ultraspeed 2 (JPK, Berlin, Germany) with silicon tips (FASTSCAN-A, drive frequency 1400 kHz, tip radius 5 nm, Bruker, Billerica, Massachusetts). Here, images were scanned over a field of view of 6 μm x 6 μm at 4096 x 4096 pixels with a scanning speed of 1.5 Hz or over a field of view of 12 μm x 12 μm at 8192 x 8192 pixels at 1.5 Hz (Fig. 1b). For each nucleosome type, four independent data sets were recorded with samples from separate nucleosome reconstitutions deposited on new muscovite mica.

### AFM image analysis

To analyze the flattened AFM images we used an analysis pipeline based in parts on the previously published, open source automated image analysis pipeline^38^. In short, image analysis consists of three steps. First, molecules are detected and classified. For molecule detection, a Gaussian filter and a background subtraction are applied to the flattened AFM images and subsequently a skeletonization^65^ – an algorithm that narrows down the objects to a one pixel wide backbone – is performed (Supplementary Fig. 1). The skeleton of the molecules is used for classification: Bare DNA has exactly two endpoints in its skeleton and nucleosomes have exactly two endpoints and two branchpoints – points with more than two neighbors (Supplementary Fig. 1). Second, a deconvolution is applied (see below). Third, the classified molecules are analyzed with respect to the structure parameters arm length, volume and opening angle for nucleosomes and length for bare DNA (Supplementary Fig. 1). Our AFM analysis code including a detailed installation guide and an example image is available on https://github.com/SKonrad-Science/AFM_nucleoprotein_readout.

### Image deconvolution

An image deconvolution is applied to the AFM images to trace nucleosomal opening angles and DNA length more accurately (see Fig. 1). Before deconvolution can be performed, the tip shape must be estimated. The estimation is done for each AFM image individually since tip shape can vary significantly even for tips of the same batch and it can change while measuring one data set over the course of several hours. To estimate the tip shape, bare DNA strands are traced without deconvolution to obtain an initial trace. Based on this initial trace a grid of 10 pixels (~15 nm) in size is filled by the height values surrounding the initial trace of the DNA strand (Supplementary Fig. 2). After repeating this process for all DNA strands in the image and averaging the intensities in the grid, a good estimate of the tip shape is available to apply the deconvolution algorithm. Here, we make use of the Richardson-Lucy deconvolution algorithm^41,42^, an iterative procedure for recovering an underlying image blurred by a known point spread function, *i.e*. the tip shape.

### AFM image simulations

To simulate nucleosome images with different levels of anti-cooperative unwrapping, an 11 nm diameter disk of uniform height together with protruding DNA arms based on the worm-like chain model at opening angles deduced from the nucleosome crystal structure (PDB 1KX5) was placed on a flat surface. The fully wrapped lengths of the short and long arm comprise 106 bp and 233 bp, respectively, and lengths were increased in 5 bp steps to simulate the individual unwrapping steps up to 35 bp. The simulated populations for each unwrapping state were chosen based on the experimentally determined populations. For simulation of anti-cooperative unwrapping, the length of only one arm was increased while keeping the length of the other arm constant. For simulation of stochastic unwrapping, the arms were randomly unwrapped in 5 bp steps up to the total amount of unwrapping simulated for each state. For example, when simulating a state of 10 bp unwrapped, possible lengths for the short arm are [106, 111, 116] bp and [233, 238, 243] bp for the long arm and were assigned randomly for each simulated nucleosome while keeping the total of 10 bp unwrapped between the two arms constant.

Consecutively, the DNA arms were dilated to their expected with of 2 nm and random noise in combination with a Gaussian filter (*σ* = 2 nm) were applied to mimic the effect of tip convolution.

## Supporting Citations

References (^66,67^) appear in the Supporting Material

