## Supplementary information for "Quantifying epigenetic modulation of nucleosome breathing by high-throughput AFM imaging"

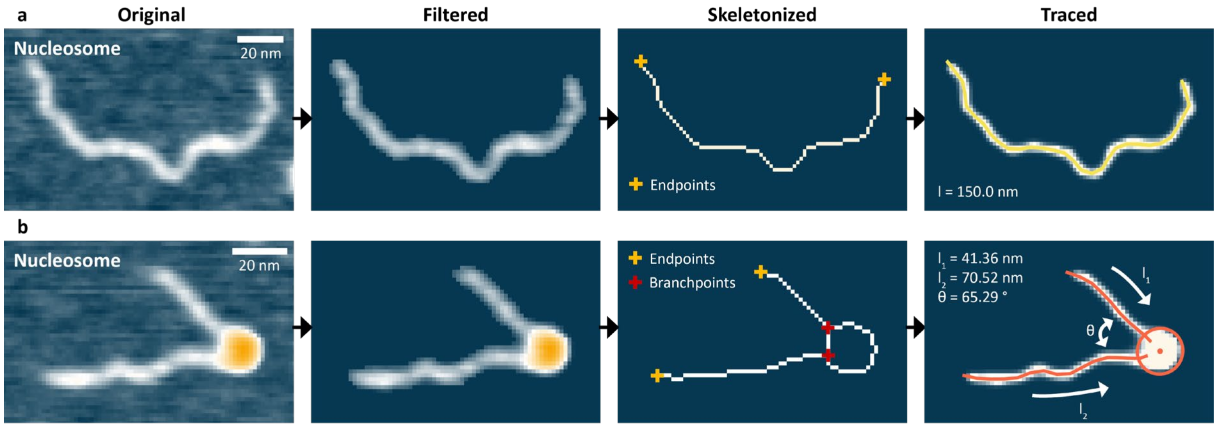


**Supplementary Figure 1 | Tracing of bare DNA and nucleosomes.**

**a,** AFM topographic image of a bare DNA strand with the different tracing steps visualized. At first, the original image is filtered by applying a Gaussian filter and removing the background with a fixed threshold value. Subsequently, the filtered image is skeletonized. The skeletonized backbone of the molecules serves as the basis for classification: whereas the skeleton of bare DNA has exactly two endpoints and no branchpoints – points that have more than two neighbors – the skeleton of nucleosomes contains exactly two endpoints and two branchpoints. Finally, the bare DNA molecule is traced with regards to its length after applying a deconvolution.

**b**, AFM topographic image of a nucleosome with the different tracing steps involved. After initial filtering, the molecule is skeletonized for classification. Finally, the nucleosome is traced with regards to the arm lengths, the opening angle and the volume.

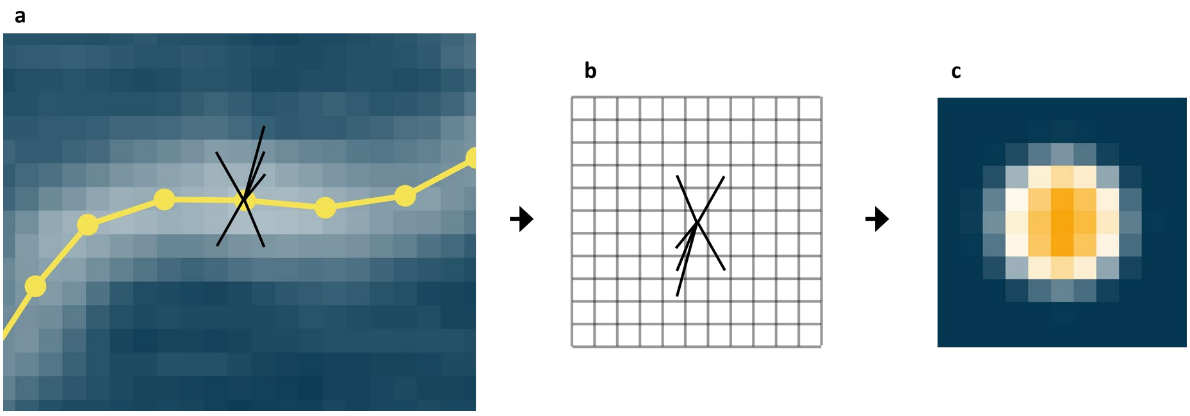


**Supplementary Figure 2 | Tip shape estimation.**

Tip shape estimation is based on analysis of the bare DNA images present in each AFM field of view.

**a,** In the first step of tip shape estimation, the vectors between the pixel centers and the nearest trace point are computed. The trace points are obtained by first tracing the DNA strand without deconvolution to get an estimate of the trace. After obtaining the tip shape and applying deconvolution, the DNA strands are traced again. All pixels that are within a maximum range of 6 nm towards the next trace point are taken into account. The amount of initial trace points was reduced for clarity in the graph. During application, the trace is approximated by a spline interpolation to provide more trace points and thus reduce error in estimating the tip shape due to a too coarse grained trace.

**b**, The vectors are then brought into an empty 2D grid and the measured height values of the pixels the vectors were obtained from in panel **a** are added to the grid. Repeating the process for multiple DNA strands (typically 100-200 DNA strands per image) will result in tens of thousands height values of vectors to fill the 2D grid with.

**c**, After adding up all height values of the vectors in the grid, the grid pixels are normalized based on the number of values added per grid pixel. The resulting distribution is an approximation of the tip shape that was used in imaging the respective image.

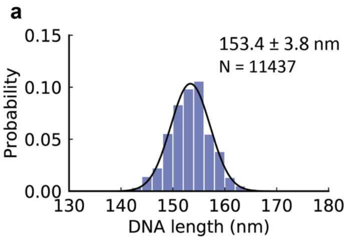


**Supplementary Figure 3 | Bare DNA lengths.**

**a,** Histogram of bare DNA lengths combined for all data sets used in this work. We find a contour length of $l_c$ = 153.4 ± 3.9 nm (mean ± std from 11437 molecules) corresponding to a length per bp of 0.316 ± 0.008 nm, in agreement with previous measurements by AFM[1,2], and solution X-ray scattering[3].

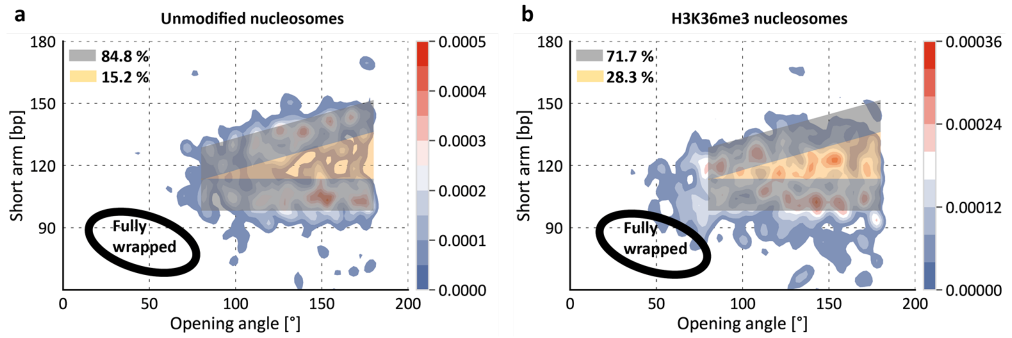


**Supplementary Figure 4 | Quantification of anti-cooperative unwrapping.**

**a,** 2D kernel density profile (bandwidth = 2.5°, 2.5 bp) of short arm length and opening angle for partially unwrapped canonical nucleosomes (N = 1035). Nucleosomes that unwrap anti-cooperatively, *i.e.* with either the long arm or the short arm unwrapping, are expected in the dark area. Nucleosomes that unwrap from both sides simultaneously are expected in the yellow area. For this particular data set of canonical nucleosomes, 84.8 % of the nucleosomes are in the regime of anti-cooperative unwrapping and 15.2 % in the regime of stochastic unwrapping. The values obtained in this analysis serve as basis for the quantification of anti-cooperativity in Fig. 4c of the main text. Per definition, the dark area makes up 75 % of the total area and the yellow area makes up 25 % of the total area. The black ellipse indicates the position of fully wrapped nucleosomes that are omitted for clarity.

**b**, 2D kernel density profile (bandwidth = 2.5°, 2.5 bp) of short arm length and opening angle for partially unwrapped H3K36me3 nucleosomes (N = 1155). 71.7 % of the nucleosomes are in the regime of anti-cooperative unwrapping and 28.3 % in the regime of stochastic unwrapping.

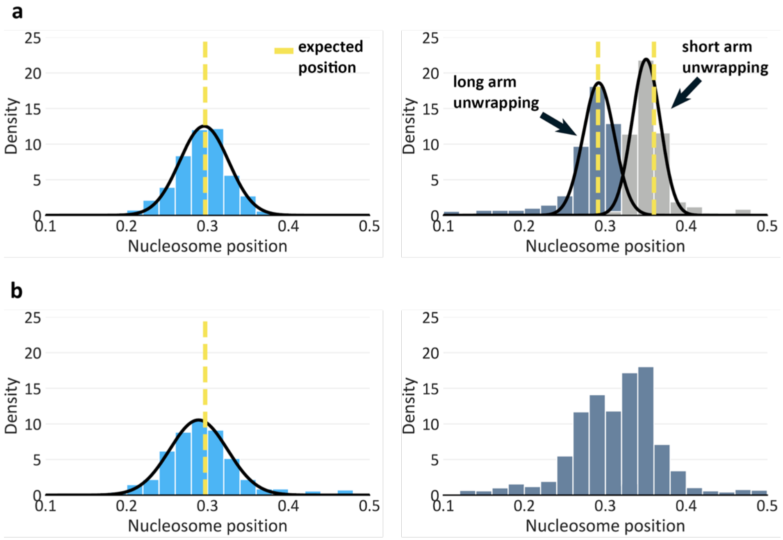


**Supplementary Figure 5 | Nucleosome positioning.**

**a,** Nucleosome positioning for a sample data set of unmodified nucleosomes (N = 1300). The nucleosome position is calculated by dividing the short arm length by the sum of short arm and long arm length. The plots show that both fully wrapped and partially unwrapped nucleosomes are positioned well at the Widom positioning sequence.

**b**, Nucleosome positioning for a sample data set of H3K36me3 nucleosomes (N = 1732). Similarly to the unmodified nucleosomes, the H3K36me3 nucleosomes are positioned well. Due to the stochastic unwrapping of H3K36me3 nucleosomes it is not possible to separate the partially unwrapped nucleosomes (right histogram) into nucleosomes that solely unwrap from the short arm and those that unwrap solely from the long arm.

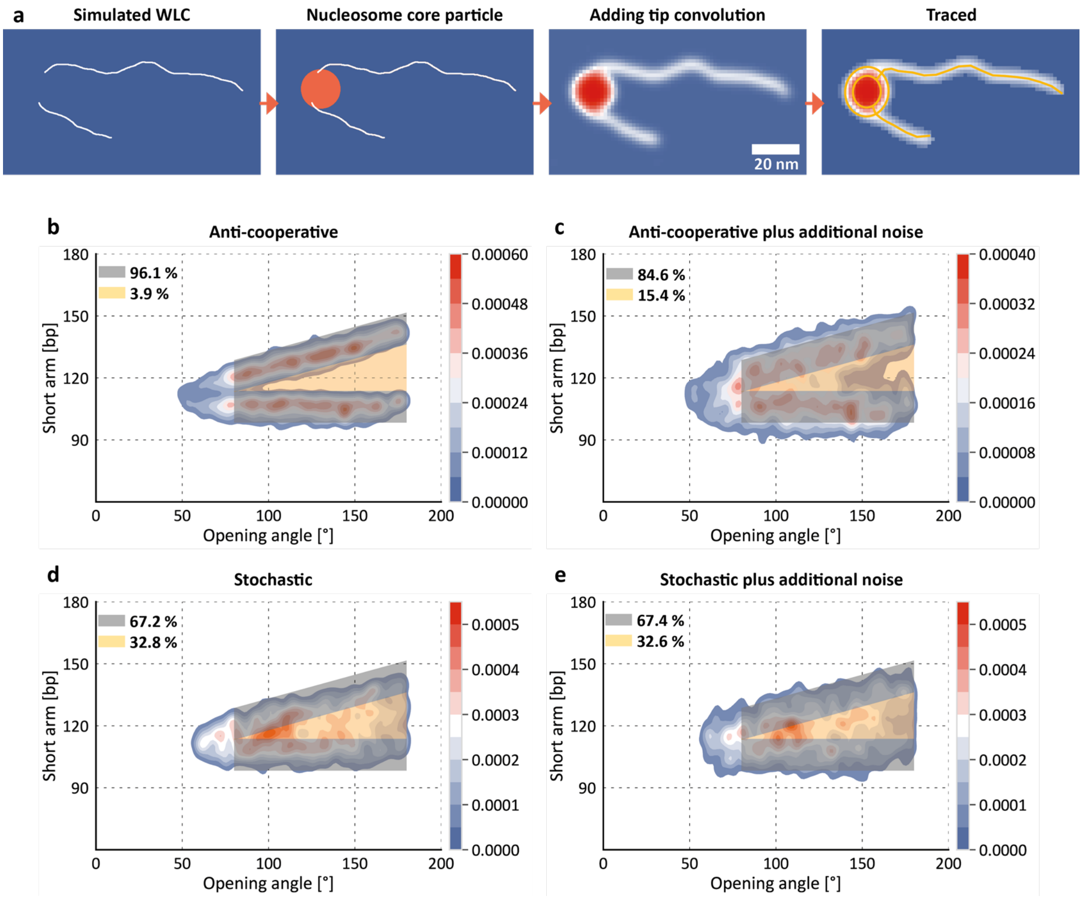


**Supplementary Figure 6 | Simulation of anti-cooperative and stochastic unwrapping.**

**a,** Simulation of nucleosomes consisted of placing a nucleosomal disk and simulating protruding DNA arms. The position and initial directionality of the protruding DNA arms was deduced from the nucleosome crystal structure (PDB 1KX5). The lengths of the DNA arms are 106 bp for the short arm and 233 bp for the long arm initially and are varied based on the state of unwrapping that is simulated (see Methods). Consecutively, the DNA was dilated to its expected width of 2 nm and a Gaussian filter was applied to mimic the effect of tip convolution. Finally, the synthetic AFM image is traced with our automated image analysis pipeline.

**b**, 2D Kernel density plot for simulated nucleosomes (N = 2072). To simulate anti-cooperative unwrapping, the length of either the short or the long arm was always kept constant (106 bp or 233 bp for the short and the long arm respectively) and the length of the other arm was increased in 5 bp steps up to a maximum unwrapping of 35 bp. Unwrapping is simulated to occur from each arm in 50 % of the cases and the sizes of the individual unwrapping populations are based on the probability for each population as experimentally measured for unmodified nucleosomes (Figure 3a).

**c**, 2D Kernel density plot for simulated nucleosomes (N = 2072). The plot comprises the same nucleosomes as shown in b. However, additional Gaussian distributed noise with $\sigma$ = 5 bp was added to the short arm length to better capture expected imaging errors that might occur during experiments.

**d**, 2D Kernel density plot for simulated nucleosomes (N = 1469). Compared to the anti-cooperative unwrapping of plots b and c, this plot shows nucleosomes that unwrap stochastically. To simulate stochastic unwrapping, the length of both arms was randomly increased in 5 bp steps for the individual unwrapping steps. For example, for simulation of states of partial unwrapping of 10 bp in total, the length of the short and long arm was randomly either increased by [+0, +10], [+5, +5] or [+10, +0] respectively. This procedure was again repeated up to a maximum unwrapping of 35 bp in total.

**e**, 2D Kernel density plot for simulated nucleosomes (N = 1469). The plot comprises the same nucleosomes as shown in d. Similar to c, additional Gaussian noise with $\sigma$ = 5 bp was added to the short arm length to better capture imaging errors that might occur during experiments.
